# Raman2RNA: Live-cell label-free prediction of single-cell RNA expression profiles by Raman microscopy

**DOI:** 10.1101/2021.11.30.470655

**Authors:** Koseki J. Kobayashi-Kirschvink, Shreya Gaddam, Taylor James-Sorenson, Emanuelle Grody, Johain R. Ounadjela, Baoliang Ge, Ke Zhang, Jeon Woong Kang, Ramnik Xavier, Peter T. C. So, Tommaso Biancalani, Jian Shu, Aviv Regev

## Abstract

Single cell RNA-Seq (scRNA-seq) and other profiling assays have opened new windows into understanding the properties, regulation, dynamics, and function of cells at unprecedented resolution and scale. However, these assays are inherently destructive, precluding us from tracking the temporal dynamics of live cells, in cell culture or whole organisms. Raman microscopy offers a unique opportunity to comprehensively report on the vibrational energy levels of molecules in a label-free and non-destructive manner at a subcellular spatial resolution, but it lacks in genetic and molecular interpretability. Here, we developed Raman2RNA (R2R), an experimental and computational framework to infer single-cell expression profiles in live cells through label-free hyperspectral Raman microscopy images and multi-modal data integration and domain translation. We used spatially resolved single-molecule RNA-FISH (smFISH) data as anchors to link scRNA-seq profiles to the paired spatial hyperspectral Raman images, and trained machine learning models to infer expression profiles from Raman spectra at the single-cell level. In reprogramming of mouse fibroblasts into induced pluripotent stem cells (iPSCs), R2R accurately (r>0.96) inferred from Raman images the expression profiles of various cell states and fates, including iPSCs, mesenchymal-epithelial transition (MET) cells, stromal cells, epithelial cells, and fibroblasts. R2R outperformed inference from brightfield images, showing the importance of spectroscopic content afforded by Raman microscopy. Raman2RNA lays a foundation for future investigations into exploring single-cell genome-wide molecular dynamics through imaging data, *in vitro* and *in vivo*.

## Main

Cellular states and functions are determined by a dynamic balance between intrinsic and extrinsic programs. Dynamic processes such as cell growth, stress responses, differentiation, and reprogramming are not determined by a single gene, but by the orchestrated temporal expression and function of multiple genes organized in programs and their interactions with other cells and the surrounding environment^1^. To understand how cells change their states in physiological and pathological conditions it is essential to decipher the dynamics of the underlying gene programs.

Despite major advances in single cell genomics and microscopy, we still cannot track live cells and tissues at the genomic level. On the one hand, single cell and spatial genomics have provided a view of gene programs and cell states at unprecedented scale and resolution^1^, but these measurement methods are destructive, and involve tissue fixation and freezing and/or cell lysis, precluding us from directly tracking the dynamics of full molecular profiles in live cells or organisms. While advanced computational methods, such as pseudo-time algorithms (*e.g*., Monocle^2^, Waddington-OT^3^) and velocity-based methods (*e.g.*, velocyto^4^, scVelo^5^), can infer dynamics from snapshots of molecular profiles, they rely on assumptions that remain challenging to verify experimentally^6^. On the other hand, fluorescent reporters can be used to monitor the dynamics of individual genes and programs within live cells, but are limited in the number of targets they can report^7^, must be chosen ahead of the experiment and often involve genetically engineered cells. Moreover, the vast majority of dyes and reporters require fixation or can interfere with nascent biochemical processes and alter the natural state of the gene of interest^7^. Therefore, it remains technically challenging to dynamically monitor the activity of a large number of genes simultaneously.

Raman microscopy opens a unique opportunity for monitoring live cells and tissues, as it collectively reports on the vibrational energy levels of molecules in a label-free and non-destructive manner at a subcellular spatial resolution, thus providing molecular fingerprints of cells^8^. Pioneering research has demonstrated that Raman microscopy can be used for characterizing cell types and cell states^8^, non-destructively diagnosing pathological specimens such as tumors^9^, characterizing the developmental states of embryos^10^, and identifying bacteria with antibiotic resistance^11^. However, the complex and high-dimensional nature of the spectra, the spectral overlaps of biomolecules such as proteins and nucleic acids, and the lack of unified computational frameworks have hindered the decomposition of the underlying molecular profiles^7, 8^.

To address this challenge and leverage the complementary strengths of Raman microscopy and scRNA-Seq, we developed Raman2RNA (R2R), an experimental and computational framework for inferring single-cell RNA expression profiles from label-free non-destructive Raman hyperspectral images (**Fig. 1**). R2R takes as input spatially resolved hyperspectral Raman images from live cells, smFISH data of selected markers from the same cells, and scRNA-seq from the same biological system. R2R then uses the smFISH data as an anchor to learn a model that links spatially resolved hyperspectral Raman images to scRNA-seq. Finally, from this model, R2R then computationally infers the anchor smFISH measurements from hyperspectral Raman images and then the single-cell expression profiles. The result is a label-free live-cell inference of single-cell expression profiles.

**Fig. 1.**
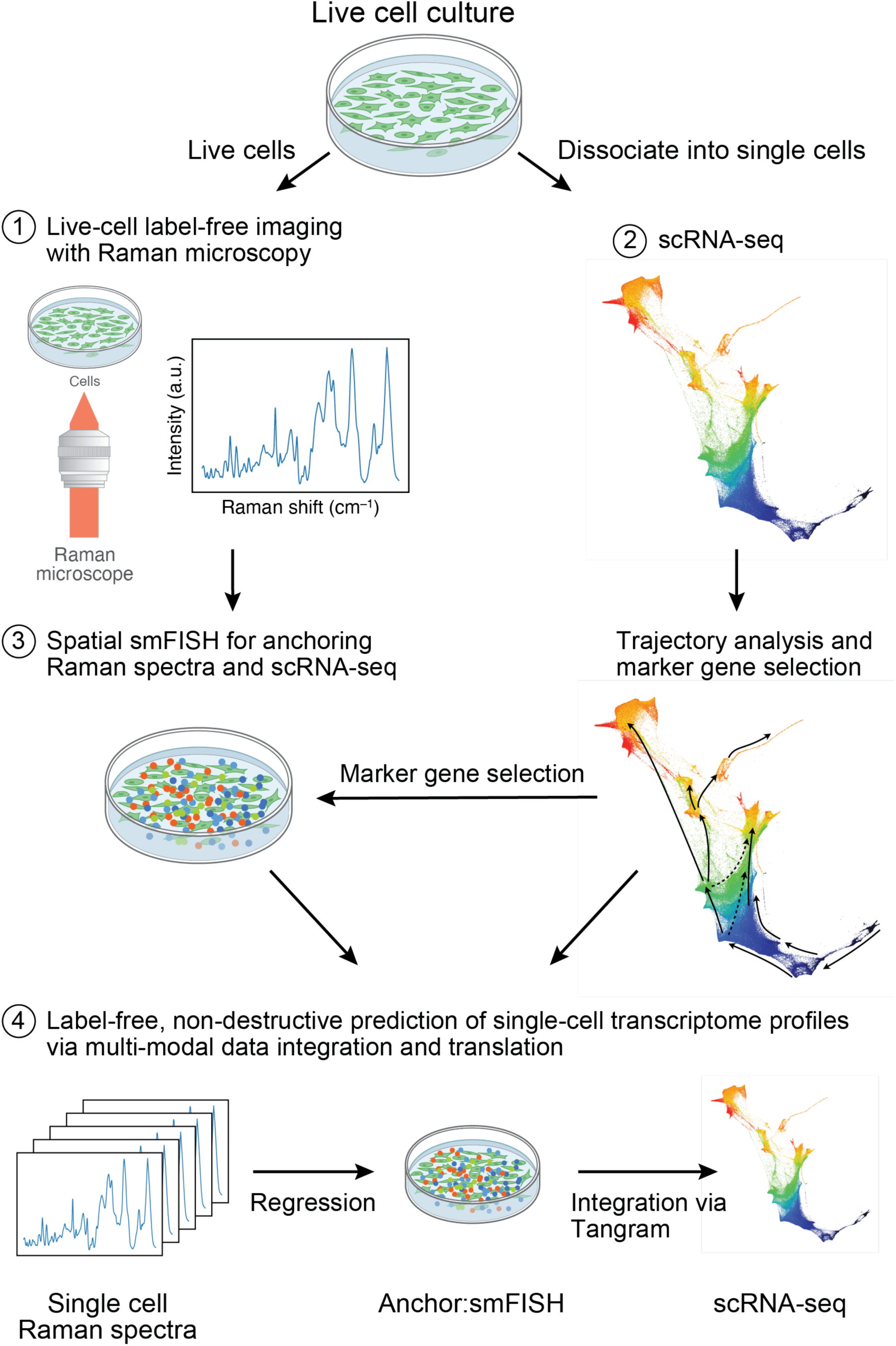
Raman2RNA. Live cells are cultured on gelatin-coated quartz glass-bottom plates (top) and Raman spectra are then measured at each pixel (at spatial sub-cellular resolution) within an image frame (1), followed by smFISH imaging in the same area (3). From parallel plates, cells are dissociated into a single cell suspension and profiled by scRNA-seq (2). scRNA-seq profiles are used to select 9 marker genes for 5 major cell clusters, and those are measured with spatial smFISH (3). Lastly, a regression model is trained (4) to predict anchor smFISH profiles from Raman spectra, followed by integration via Tangram^16^ to predict whole single-cell transcriptome profiles from smFISH profiles.

To facilitate data acquisition, we developed a high-throughput multi-modal spontaneous Raman microscope that enables automated acquisition of Raman spectra, brightfield, and fluorescent images. In particular, we integrated Raman microscopy optics to a fluorescence microscope, where high-speed galvo mirrors and motorized stages were combined to achieve a large field of view (FOV) scanning, and where dedicated electronics automate measurements across multiple modalities (**Extended Data Fig. 1-2, Methods**).

We first demonstrated that R2R can infer profiles of two distinct cell types: mouse induced pluripotent stem cells (iPSCs) expressing an endogenous *Oct4*-GFP reporter and mouse fibroblasts^12^. To this end, we mixed the cells in equal proportions, plated them in a gelatin-coated quartz glass-bottom Petri dish, and performed live-cell Raman imaging, along with fluorescent imaging of live-cell nucleus staining dye (Hoechst 33342) for cell segmentation and image registration, and an iPSC marker gene, *Oct4*-GFP (**Fig. 2a**). The excitation wavelength for our Raman microscope (785 nm) was distant enough from the GFP Stokes shift emission, such that there was no interference with the cellular Raman spectra (**Extended Data Fig. 3**). Furthermore, there was no notable photo-toxicity induced in the cells. After Raman and fluorescence imaging, we fixed and permeabilized the cells and performed smFISH (with hybridization chain reaction (HCR^13^), **Methods**) of marker genes for mouse iPSCs (*Nanog*) and fibroblasts (*Col1a1*). We registered the nuclei stains, GFP images, HCR images, and Raman images through either polystyrene control bead images or reference points marked under the glass bottom dishes (**Extended Data Fig. 4, Methods**).

**Fig. 2.**
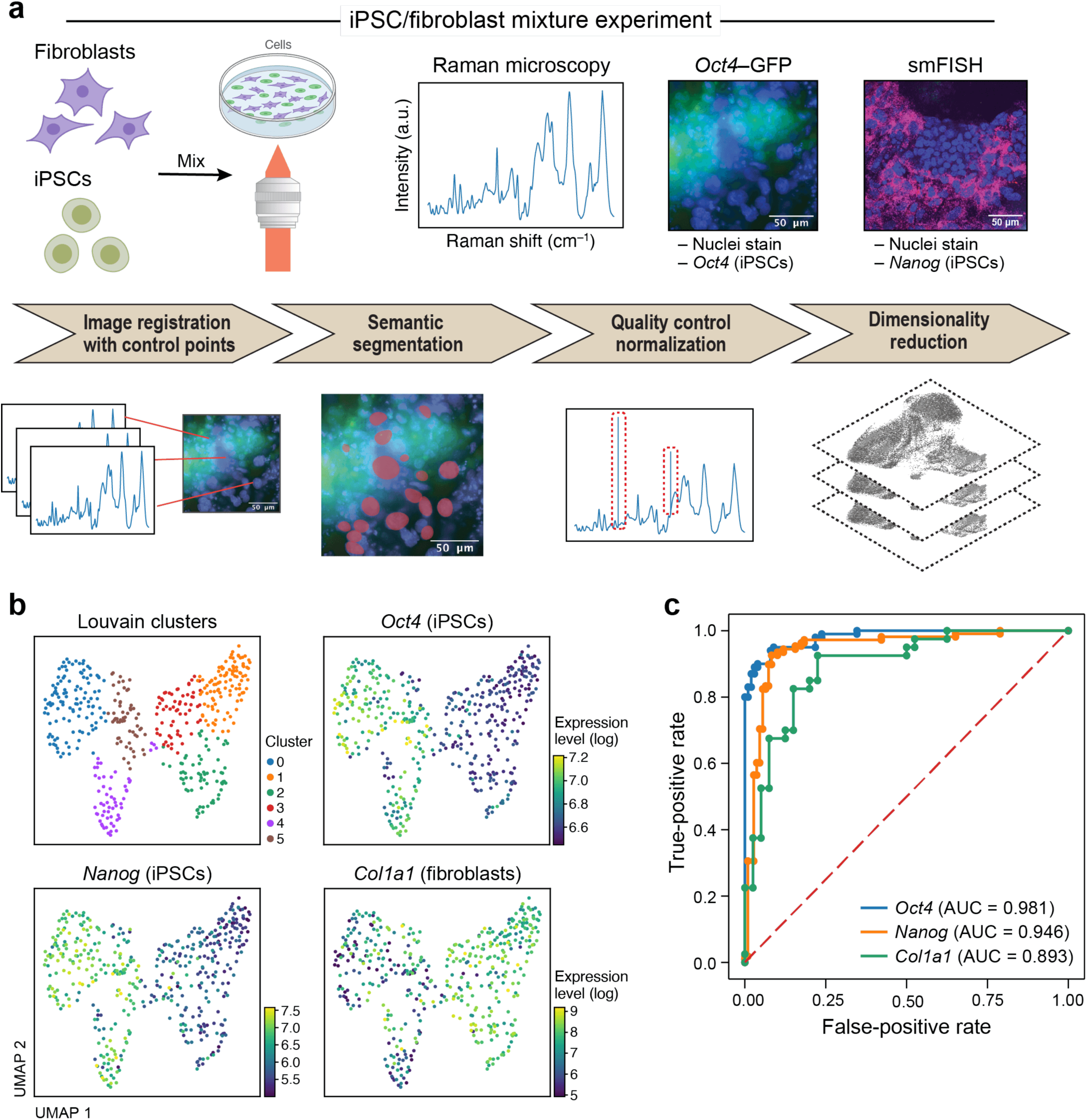
Raman2RNA accurately distinguishes cell types and predicts binary expression of marker genes in a mixture of mouse fibroblasts and iPSCs. **a.** Overview. Top: Experimental procedures. Mouse fibroblasts and iPSCs were mixed 1:1 and plated on glass-bottom plates, followed by Raman imaging of live cells, nuclei staining and measurement of endogenous *Oct4*-GFP (iPSC marker) reporter) by fluorescence imaging, and cell fixation and processing for smFISH with DAPI and probes for *Nanog* (iPSCs, magenta) and *Col1a1* (fibroblasts). Bottom: Preprocessing and analysis. From left: Image registration with control points (**Methods**), was followed by semantic cell segmentation, outlier removal/normalization and dimensionality reduction. **b.** Raman2RNA distinguishes cell states from Raman spectra. 2D UMAP embedding of single-cell Raman spectra (dots) colored by Louvain clustering labels (top left) or smFISH measured expression of *Oct4* (top right), *Nanog* (bottom left) and *Col1a1* (bottom right). **c.** Raman2RNA accurately predicts binary (on/off) expression of marker genes. Receiver operating characteristic (ROC) plots and area under the curve (AUC) obtained by classifying the ‘on’ and ‘off’ states of *Oct4* (blue), *Nanog* (orange) and *Col1a1* (green).

The Raman spectra distinguished the two cell populations in a manner congruent with the expression of their respective reporter (measured live or by smFISH in the same cells), as reflected by a low-dimensional embedding of hyperspectral Raman data (**Fig. 2b**). Specifically, we focused on the fingerprint region of Raman spectra (600-1800 cm^-1^, 930 of the 1,340 features in a Raman spectrum), where most of the signatures from various key biomolecules, such as proteins, nucleic acids, and metabolites, lie^8^. After basic preprocessing, including cosmic-ray and background removal and normalization, we aggregated Raman spectra that are confined to the nuclei, obtaining a 930-dimensional Raman spectroscopic representation for each cell’s nucleus. We then visualized these Raman profiles in an embedding in two dimensions using Uniform Manifold Approximation and Projection (UMAP)^14^ and labeled cells with the gene expression levels that were concurrently measured by either an *Oct4*-GFP reporter or smFISH (**Fig. 2b**). The cells separated clearly in their Raman profiles in a manner consistent with their gene expression characteristics, forming two main subsets in the embedding, one with cells with high *Oct4* and *Nanog* expression (iPSCs markers) and another with cells with relatively high *Col1a1* expression (fibroblasts marker), indicating that Raman spectra reflect cell-intrinsic expression differences (**Fig. 2b**).

We further successfully trained a classifier to classify the ‘on’ or ‘off’ expression states of *Oct4*, *Nanog* and *Col1a1* in each cell based on its Raman profile (**Methods**). We trained a logistic regression classifier with 50% of the data and held out 50% for testing. We predicted *Oct4* and *Nanog* expression states with high accuracy on the held-out test data (area under the receiver operating characteristic curve (AUROC) = 0.98 and 0.95, respectively; **Fig. 2c**), indicating that expression of iPSC markers can be predicted confidently from Raman spectra of live, label-free cells. We also successfully classified the expression state of the fibroblast marker *Col1a1* (AUROC = 0.87; **Fig. 2c**), albeit with lower confidence, which is consistent with the lower contrast in *Col1a1* expression (**Fig. 2b**) between iPSC (*Oct4*+ or *Nanog*+ cells) *vs*. non-iPSCs, compared to *Oct4* or *Nanog*. Most misclassifications occurred when the ground truth expression levels were near the threshold of the classifier, showing that misclassifications were likely due to the uncertainty in the ground truth expression level (**Extended Data Fig. 5**).

Next, we asked if the Raman images could predict entire expression profiles non-destructively at single-cell resolution. To this end, we aimed to reconstruct scRNA-seq profiles from Raman images by multi-modal data integration and translation, using multiplex smFISH data to anchor between the Raman images and scRNA-seq profiles (**Fig. 3a**). As a test case, we focused on the mouse iPSC reprogramming model system, where we have previously generated ∼250,000 scRNA-seq profiles at ½ day intervals throughout an 18 day, 36 time point time course of reprogramming^3^ (**Methods**). We used Waddington-OT^3^ (WOT) to select from the scRNA-seq profiles nine anchor genes that represent diverse cell types that emerge during reprogramming (iPSCs: *Nanog*, *Utf1* and *Epcam*; MET and neural: *Nnat* and *Fabp7*; epithelial: *Krt7* and *Peg10*; stromal: *Bgn* and *Col1a1*; **Methods**). We performed live-cell Raman imaging from day 8 of reprogramming, in which distinct cell types begin to emerge^3^, up to day 14.5, at half-day intervals, totaling 14 time points (**Methods**). We imaged ∼500 cells per plate at 1µm spatial resolution. Finally, we fixed cells immediately after each Raman imaging time point followed by smFISH on the 9 anchor genes (**Methods**).

**Fig. 3.**
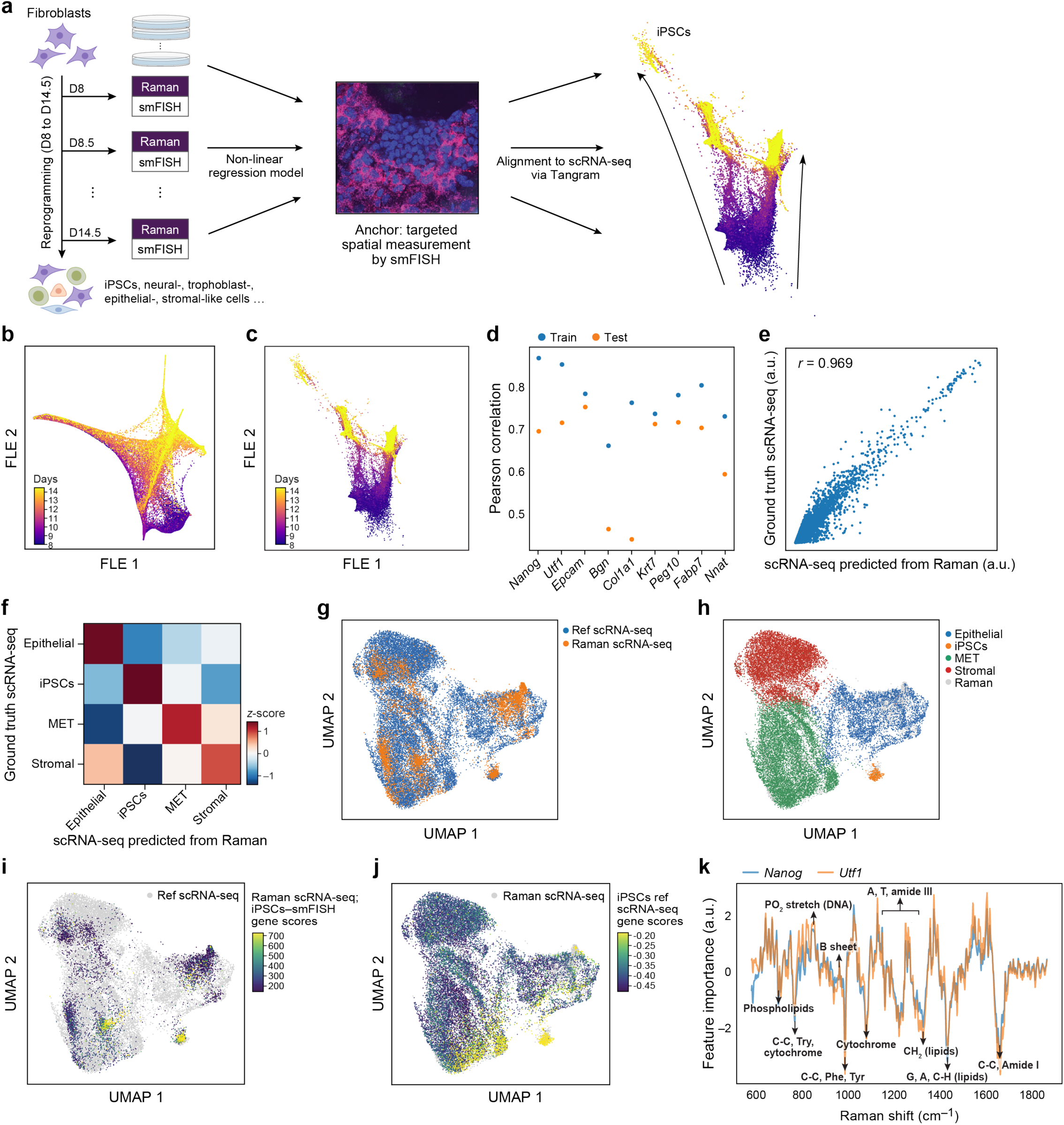
Raman2RNA predicts single-cell RNA profiles across cell types during reprogramming of mouse fibroblasts to iPSCs. **a.** Approach overview. From left: Mouse fibroblasts were reprogrammed into induced pluripotent stem cells (iPSCs) over the course of 14.5 days (‘D’), and, at half-day intervals from days 8 to 14.5, spatial Raman spectra, smFISH for nine anchor genes, and nuclei stain by fluorescence imaging were measured for each plate. Machine learning and multi-modal data integration methods (Catboost and Tangram) were used to predict single-cell RNA-seq profiles from Raman spectra using smFISH as anchor. **b,c.** Low dimensionality embedding of single-cell Raman spectra captures progress in reprogramming. Force-directed layout embedding (FLE) of Raman spectra (b, dots) or scRNA-seq (c, dots) colored by days of measurement (colorbar). **d.** Correct prediction of smFISH anchors from Raman spectra. Pearson correlation coefficient (*y* axis) between measured (smFISH) and Raman-predicted levels for each smFISH anchor (*x* axis) in leave-one-out cross-validation where 8 out of 9 smFISH anchor genes were used for training, and the left-out gene was predicted. **e.f.** Raman2RNA accurately predicts pseudo-bulk expression profiles of major cell types. **e.** scRNA-seq measured (y axis) and R2R-predicted (x axis) for each gene (dot) in pseudo-bulk RNA profiles averaged across iPSCs. **f.** Pair-wise correlation (color bar) between Raman-predicted and scRNA-seq measured pseudo-bulk profiles in each cell types (rows, columns). **g-j.** Co-embedding highlights agreement between real and R2R inferred single cell profiles. UMAP co-embedding of Raman predicted RNA profiles and measured scRNA-seq profiles (dots) colored by data source (**g,** Raman predicted in orange; measured scRNA-seq in blue), cell type annotations (**h**) or by iPSC gene signature scores (calculated by averaging expression of genes *Nanog* and *Utf1*, and subtracting the average of a randomly selected set of reference genes; **Methods**) of Raman-predicted profiles (**i**) or of real scRNA-seq (**j**). **k.** Feature importance scores of Raman spectra in predicting expression profiles. Feature scores for iPSC related marker genes (y axis) along the Raman spectrum (x axis). Known Raman peaks^18^ were annotated.

Strikingly, a low dimensional representation of the Raman profiles showed that they encoded similar temporal dynamics to those observed with scRNA-seq during reprogramming (**Fig. 3b,c, Extended Data Fig. 6**), indicating that they may qualitatively mirror scRNA-seq.

Integrating Raman and scRNA-seq profiles (**Methods**), R2R then learned a model that can infer an scRNA-seq profile for each Raman imaged cell, by first predicting smFISH anchors from the Raman profiles using Catboost^15^ (**Methods**) and then using our Tangram^16^ method to map from the anchors to full scRNA-seq profiles (**Fig. 1**, **Fig. 3d-f**). In the first step, we averaged the smFISH signal within a nucleus to represent a single nucleus’s expression level. As we conducted smFISH of 9 genes, the result was a 9-dimensional smFISH profile for each single nucleus. Then, Raman profiles were translated to these 9-dimensional profiles with Catboost^15^, a non-linear regression model, using 50% of the Raman and smFISH profiles as training data.

In the second step, we mapped these anchor smFISH profiles to full scRNA-seq profiles using Tangram, yielding well-predicted single cell RNA profiles, as supported by several lines of evidence. First, we performed leave-one-out cross-validation (LOOCV) analysis, in which we used eight out of the nine anchor genes to integrate Raman with scRNA-seq, and compared the predicted expression of the remaining genes to its smFISH measurements. The predicted left-out genes based on scRNA-seq showed a significant correlation with the measured smFISH expression for any left-out gene (Pearson *r*∼0.7, *p*-value<10^-100^, **Fig. 3d**). Notably, when we analogously applied a modified U-net^17^ to infer smFISH profiles from brightfield (**Extended Data Fig. 15, Methods**), we observed a poor, near-random prediction of expression profiles for all 9 genes in leave-one-out cross-validation (*r*<0.15), indicating that, unlike Raman spectra, brightfield z-stack images either do not have the necessary information to infer expression profiles, or require more data. Second, we compared the real (scRNA-seq measured) and R2R predicted expression profiles averaged across cells of the same cell type (“pseudobulk” for each of iPSCs, epithelial cells, stromal cells, and MET). Here, we obtained the “ground truth” cell types of the R2R profiles by transferring scRNA-seq annotations to the matching smFISH profiles using Tangram’s label transfer function. Then, based on the labels, we averaged R2R’s predicted profiles across the cells of a single cell type. The two profiles (R2R-inferred and scRNA-seq pseudo-bulk per cell type) showed high correlations (Pearson’s *r*>0.96) (**Fig. 3e,f**, **Extended Data Fig. 7**), demonstrating the accuracy of R2R at the cell type level. Furthermore, projecting the R2R predicted profiles of each cell onto an embedding learned from the real scRNA-seq shows that the predicted profiles span the key cell types as captured in real profiles (**Fig. 3g-j, Extended Data Fig. 8-12**). We note that the predicted profiles had lower variance compared to real scRNA-seq. As this is observed even when co-embedding only smFISH and scRNA-seq measurements (with no Raman data or projection, **Extended Data Fig. 13**), we believe it mostly reflects the limited number and domain maladaptation of the smFISH anchor genes used for integration. Given the similarity of the separate embeddings of Raman and scRNA-seq profiles, future studies without anchors could address this.

Lastly, we calculated feature importance scores in R2R predictions (**Methods**) and identified Raman spectral features correlated with expression levels (**Fig. 3k**, **Extended Data Fig. 14**). For example, Raman bands at approximately 752cm^-1^ (C-C, Try, cytochrome), 1004 cm^-1^ (CC, Phe, Tyr), and 1445 cm^-1^ (CH_2_, lipids) contributed to predicting iPSCs-related expression profiles, which is consistent with previous research that employed single cell Raman spectra to identify mouse embryonic stem cells (ESCs)^18^ (**Fig. 3k**). The contributions of these bands were either suppressed or increased for other cell types, such as stromal or epithelial cells (**Extended Data Fig. 14**).

In conclusion, we reported R2R, a label-free non-destructive framework for inferring expression profiles at single-cell resolution from Raman spectra of live cells, by integrating Raman hyperspectral images with scRNA-seq data through paired smFISH measurements and multi-modal data integration and translation. We inferred single-cell expression profiles with high accuracy, based on both averages within cell types and co-embeddings of individual profiles. We further showed that predictions using brightfield z-stacks had poor performance, indicating the importance of Raman microscopy for predicting expression profiles.

R2R can be further developed in several ways. First, the throughput of single-cell Raman microscopy is still limited. In this pilot study, we profiled ∼6,000 cells in total. By using emerging vibrational spectroscopy techniques, such as Stimulated Raman Scattering microscopy^19^ or photo-thermal microscopy^20, 21^, we envision increasing throughput by several orders of magnitude, to match the throughput of massively parallel single cell genomics. Second, because molecular circuits and gene regulation are structured, with strong co-variation in gene expression profiles across cells, we can leverage the advances in computational microscopy to infer high-resolution data from low-resolution data, such as by using compressed sensing, to further increase throughput^22^. Third, increasing the number of anchor genes (*e.g.*, by seqFISH^23^, merFISH^24^, STARmap^25^, or ExSeq^26^) can increase our prediction accuracy and capture more single-cell variance. Additionally, with single-cell multi-omics, we can project other modalities, such as scATAC-seq from Raman spectra. Finally, given the similarity in the overall independent embedding of Raman and scRNA-seq profiles, we expect computational methods such as multi-domain translation^27^ to allow mapping between Raman spectra and molecular profiles without measuring any anchors *in situ*. Overall, with further advances in single-cell genomics, imaging, and machine learning, Raman2RNA could allow us to non-destructively infer omics profiles at scale *in vitro*, and possibly *in vivo* in living organisms.

## Materials and Methods

### Mouse fibroblast reprogramming

OKSM secondary mouse embryonic fibroblasts (MEFs) were derived from E13.5 female embryos with a mixed B6;129 background. The cell line used in this study was homozygous for ROSA26-M2rtTA, homozygous for a polycistronic cassette carrying *Oct4*, *Klf4*, *Sox2*, and *Myc* at the *Col1a1* 3’ end, and homozygous for an EGFP reporter under the control of the *Oct4* promoter. Briefly, MEFs were isolated from E13.5 embryos from timed-matings by removing the head, limbs, and internal organs under a dissecting microscope. The remaining tissue was finely minced using scalpels and dissociated by incubation at 37°C for 10 minutes in trypsin-EDTA (ThermoFisher Scientific). Dissociated cells were then plated in MEF medium containing DMEM (ThermoFisher Scientific), supplemented with 10% fetal bovine serum (GE Healthcare Life Sciences), non-essential amino acids (ThermoFisher Scientific), and GlutaMAX (ThermoFisher Scientific). MEFs were cultured at 37°C and 4% CO_2_ and passaged until confluent. All procedures, including maintenance of animals, were performed according to a mouse protocol (2006N000104) approved by the MGH Subcommittee on Research Animal Care^3^.

For the reprogramming assay, 50,000 low passage MEFs (no greater than 3-4 passages from isolation) were seeded in 14 3.5cm quartz glass-bottom Petri dishes (Waken B Tech) coated with gelatin. These cells were cultured at 37°C and 5% CO_2_ in reprogramming medium containing KnockOut DMEM (GIBCO), 10% knockout serum replacement (KSR, GIBCO), 10% fetal bovine serum (FBS, GIBCO), 1% GlutaMAX (Invitrogen), 1% nonessential amino acids (NEAA, Invitrogen), 0.055 mM 2-mercaptoethanol (Sigma), 1% penicillin-streptomycin (Invitrogen) and 1,000 U/ml leukemia inhibitory factor (LIF, Millipore). Day 0 medium was supplemented with 2 mg/mL doxycycline Phase-1 (Dox) to induce the polycistronic OKSM expression cassette. The medium was refreshed every other day. On day 8, doxycycline was withdrawn. Fresh medium was added every other day until the final time point on day 14. One plate was taken every 0.5 days after day 8 (D8-D14.5) for Raman imaging and fixed with 4% formaldehyde immediately after for HCR.

### High-throughput multi-modal Raman microscope

Due to the lack of commercial systems, we developed an automated high-throughput multi-modal microscope capable of multi-position and multi-timepoint fluorescence imaging and point scanning Raman microscopy (**Extended Data Fig. 1**). A 749 nm short-pass filter was placed to separate brightfield and fluorescence from Raman scattering signal, and the fluorescence and Raman imaging modes were switched by swapping dichroic filters with auto-turrets. To realize a high-throughput Raman measurement, galvo mirror-based point scanning and stage scanning was combined to acquire each FOV and multiple different FOVs, respectively.

To realize this in an automated fashion, a MATLAB (2020b) script that communicates with Micro-manager^28^, a digital acquisition (DAQ) board, and Raman scattering detector (Princeton Instruments, PIXIS 100BR eXcelon) was written (**Extended Data Fig. 2**). A 2D point scan Raman imaging sequence was regarded as a dummy image acquisition in Micro-manager, during which the script communicated via the DAQ board with 1. the detector to read out a spectrum, 2. the mirror to update the mirror angles, and 3. shutters to control laser exposure. All communications were realized using transistor-transistor logic (TTL) signaling. Updating of the galvo mirror angles was conducted during the readout of the detector. While the script ran in the background, Micro-manager initiated a multi-dimensional acquisition consisting of brightfield, DAPI, GFP, and dummy Raman channel at multiple positions and z-stacks.

An Olympus IX83 fluorescence microscope body was integrated with a 785 nm Raman excitation laser coupled to the backport, where the short-pass filter deflected the excitation to the sample through an Olympus UPLSAPO 60X NA 1.2 water immersion objective. The backscattered light was collimated through the same objective and collected with a 50 μm core multi-mode fiber, which was then sent to the spectrograph (Holospec f/1.8i 785 nm model) and detector. The fluorescence and brightfield channels were imaged by the Orca Flash 4.0 v2 sCMOS camera from Hamamatsu Photonics. The exposure time for each point in the Raman measurement was 20 msec, and laser power at the sample plane was 212 mW. Each FOV was 100×100 pixels, with each pixel corresponding to about 1 μm. The laser source was a 785 nm Ti-Sapphire laser cavity coupled to a 532 nm pump laser operating at 4.7W.

The time to acquire Raman hyperspectral images was roughly 8 minutes per FOV. With 8 minutes, it is unrealistic to image an entire glass-bottom plate. Therefore, we visually chose representative FOVs that cover all representative cell types including iPSC-like, epithelial-like, stromal-like and MET cells. 20 FOVs were chosen for each plate, where roughly 15 FOVs were from the boundaries of colonies, five from non-colonies, and one from non-cells to use for background correction.

Due to the extended Raman imaging time, evaporation of the immersion water was no longer negligible. Therefore, we developed an automated water immersion feeder using syringe pumps and syringe needles glued to the tip of the objective lens. Here, water was supplied at a flow rate of 1 µL/min.

### iPSC and MEF mixture experiment

Low passage iPSCs were first cultured in N2B27 2i media containing 3 mM CHIR99021, 1 mM PD0325901, and LIF. On the day of the experiment, 750,000 iPSCs and 750,000 MEFs were plated on the same gelatin-coated 3.5cm quartz glass-bottom Petri dish. Cells were plated in the same reprogramming medium as previously described (with Dox) with the exception of utilizing DMEM without phenol red (Gibco) instead of KnockOut DMEM. 6 hours after plating, the quartz dishes were taken for Raman imaging and fixed with 4% formaldehyde immediately after for HCR.

### Anchor gene selection by Waddington-OT

To select anchor genes for connecting spatial information to the full transcriptome data, Waddington-OT (WOT)^3^, a probabilistic time-lapse algorithm that can reconstruct developmental trajectories, was used. We applied WOT to mouse fibroblast reprogramming scRNA-seq data collected at matching time-points and culture condition (day 8-14.5 at ½ day intervals)^3^. For each cell fate, we calculated the transition probabilities of each cell and selected the top 10 percentile cells per time point (**Extended Data Fig. 6**). Based on this, we ran the *FindMarker* function in Seurat^29^ to find genes differentially expressed in these cell subsets per time point. Through this approach, we chose two genes per cell type that are both found by Seurat and commonly used for these cell types (iPSCs: *Nanog*, *Utf1*; epithelial: *Krt7*, *Peg10*; stromal: *Bgn*, *Col1a1*; MET and neural: *Fabp7*, *Nnat*), along with one gene that is an early marker of iPSCs, *Epcam*.

### smRNA-FISH by hybridization chain reaction (HCR)

Fixed samples were prepared for imaging using the HCR v3.0 protocol for mammalian cells on a chambered slide, incubating at the amplification step for 45 minutes in the dark at room temperature. Three probes with amplifiers conjugated to fluorophores Alexa Fluor 488, Alexa Fluor 546, and Alexa Fluor 647 were used. Samples were stained with DAPI prior to imaging. After imaging, probes were stripped from samples by washing samples once for 5 minutes in 80% formamide at room temperature and then incubating three times for 30 minutes in 80% formamide at 37°C. Samples were washed once more with 80% formamide, then once with PBS, and reprobed with another panel of probes for subsequent imaging.

### Image registration of Raman hyperspectral images and fluorescence/smFISH images

Brightfield and fluorescence channels including DAPI and GFP, along with corresponding Raman images, were registered by using 5 μm polystyrene beads deposited on quartz glass-bottom Petri dishes (SF-S-D12, Waken B Tech) for calibration. The brightfield and fluorescence images of the beads were then registered by the scale-invariant template matching algorithm of the OpenCV (https://github.com/opencv/opencv) *matchTemplate* function followed by manual correction. For the registration of smFISH and Raman images, four marks inscribed under the glass-bottom Petri dishes were used as reference points (**Extended Data Fig. 4**). As the Petri dishes are temporarily removed from the Raman microscope after imaging to do smFISH measurements, the dishes cannot be placed back at the same exact location on the microscope. Therefore, the coordinates of these reference points were measured along with the different FOVs. When the dishes were placed again after smFISH measurements, the reference mark coordinates were measured, and an affine mapping was constructed to calculate the new FOV coordinates. Lastly, as smFISH consisted of 3 rounds of hybridization and imaging, the following steps were performed to register images across different rounds with a custom MATLAB script:

1. Maximum intensity projection of nuclei stain and RNA images
2. Automatic registration of round 1 images to rounds 2 and 3 based on nuclei stain images and MATLAB function *imregtform*. First, initial registration transformation functions were obtained with a similarity transformation model passing the ‘multimodal’ configuration. Then, those transformations were used as the initial conditions for an affine model-based registration with the *imregtform* function. Finally, this affine mapping transformation was applied to all the smFISH (RNA) images.
3. Use the protocol in (2) to register nuclei stain images obtained from the multimodal Raman microscope and the 1^st^ round of images used for smFISH. Then, apply the transformation to the remaining 2^nd^ and 3^rd^ rounds.
4. Manually remove registration outliers in (3).

Fibroblast cells were mobile during the 2-class mixture experiment so that by the time Raman imaging finished, cells had moved far enough from their original position that the above semi-automated strategy could not be applied. Thus, we manually identified cells present in both nuclei stain images before and after the Raman imaging.

### Hyperspectral Raman image processing

Each raw Raman spectrum has 1,340 channels. Of those channels, we extracted the fingerprint region (600-1800 cm^-1^), which resulted in a total of 930 channels per spectrum. Thus, each FOV is a 100×100×930 hyperspectral image. The hyperspectral images were then preprocessed by a python script as follows:

1. Cosmic ray removal. Cosmic rays were detected by subtracting the median filtered spectra from the raw spectra, and any feature above 5 was classified as an outlier and replaced with the median value. The kernel window size for the median filter was 7.
2. Autofluorescence removal. The *baseline* function in *rampy* (https://github.com/charlesll/rampy), a python package for Raman spectral preprocessing, was used with the alternating least squares algorithm *‘als’*.
3. Savitzky-Golay smoothing. The *scipy.signal.savgol_filter* function was used with window size 5 and polynomial order 3.
4. Averaging spectra at the single-cell level. Nuclei stain images were segmented using *NucleAIzer* (https://github.com/spreka/biomagdsb) and averaged pixel-level spectra that fall within each nucleus.
5. Spectra standardization. Spectra were standardized to a mean of 0 and a standard deviation of 1.

### Inferring anchor smFISH from Raman spectra or brightfield z-stacks

For the two-class mixture and reprogramming experiment, we trained a decision tree-based non-linear regression, *Catboost*^15^, to predict the ‘on’ or ‘off’ expression states for each anchor gene from Raman spectra. We used 80% of the data as training and the remaining 20% as test data. The early stopping parameter was set to 5.

For the brightfield z-stack to smFISH inference, we applied deep learning to the whole image level. We trained a modified U-net with skip connections and residual blocks to estimate the corresponding smFISH image^17^. Due to the small size of the available training dataset, we augmented the data by rotation and flipping. Furthermore, a subsample of each brightfield image was taken due to memory constraints (50×50 pixel region). Training was carried out on an NVIDIA Tesla P100 GPU, the number of epochs was 100, the learning rate was 0.01, and the batch size was 400. For each smFISH prediction, we chose the epoch that gave the best validation score.

### Inferring expression profiles from Raman images

To infer expression profiles from Raman images, we used Tangram^16^. Tangram enables the alignment of spatial measurements of a small number of genes to scRNA-seq measurements. After using Catboost to infer anchor expression levels from Raman profiles, we aligned the inferred expression levels to scRNA-seq profiles using the *map_cells_to_space* function (learning_rate=0.1, num_epochs=1000) on an Nvidia Tesla P100 GPU, followed by the *project_genes* function in Tangram.

When comparing different pseudo-bulk transcriptome predictions with the real scRNA-seq data, we first transferred labels of annotated scRNA-seq profiles to the ground truth smFISH profiles using Tangram’s label transfer function *project_cell_annotations*. Then, the average expression profiles across cells of a cell type were calculated by referring to the transferred labels and compared with those from the real scRNA-seq data^3^.

### Dimensionality reduction, embedding and projection

For dimension reduction and visualization of Raman and scRNA-seq profiles, we performed forced layout embedding (FLE) using the *Pegasus* pipeline (https://github.com/klarman-cell-observatory/pegasus). First, we performed principal component analysis on both Raman and scRNA-seq profiles independently, calculated diffusion maps on the top 100 principal components, and performed an approximated FLE graph using Deep Learning by *pegasus.net_fle* with default parameters.

To project Raman profiles to a scRNA-seq embedding, we calculated a k-nearest neighbor graph (*k*-NN, *k*=15) on the scRNA-seq top 50 principal components with the cosine metric, and UMAP with the *scanpy.tl.umap* function in Scanpy^30^ version 1.7.2 with default parameters. Then, the Raman predicted expression profiles were projected on to the scRNA-seq UMAP embedding by *scanpy.tl.ingest* using k-NN as the labeling method and default parameters.

### Feature importance analysis

To evaluate the contributions of Raman spectral features to expression profile prediction, we used the *get_feature_importance* function in Catboost with default parameters. As the dimensions of Raman spectra were reduced by PCA prior to Catboost, feature importance scores were calculated for each principal component, and the weighted linear combination of the Raman PCA eigen vectors with feature scores as the weight were calculated to obtain the full spectrum.

## Author contributions

KJKK, JS, TB and AR conceived the research and developed the methodology. JS, TB and AR funded and supervised research. KJKK, JS, JO performed reprogramming experiments. KJKK developed the multi-modal Raman microscope and control software with supervision from JWK and PS. KJKK, EG, and KZ performed smFISH. KJKK, SG, TJS, and TB developed the Raman spectral preprocessing and classification pipeline. KJKK developed the image registration pipeline, and performed Waddington-OT, Tangram and feature importance analysis. KJKK and BG performed U-net. KJKK, JS, and AR wrote the manuscript with input from all the authors.

## Acknowledgements

KJKK was supported by the Japan Society for the Promotion of Science Postdoctoral Fellowship for Overseas Researchers, and the Naito Foundation Overseas Postdoctoral Fellowship. BG was supported by the MathWorks Fellowship. JS was supported by the Helen Hay Whitney Foundation and NIH Pathway to Independence Award (1K99HD096049-01, 5K99HD096049-02, 4R00HD096049-03), and funds from the Broad Institute of MIT and Harvard and Massachusetts General Hospital. This research was funded by NIH National Institute of Biomedical Imaging and Bioengineering, grant P41EB015871 (JWK, PS), NIH grant U19 MH114821 (TB), HubMap UH3CA246632 (TB), and HHMI and the Klarman Cell Observatory (AR). AR was a Howard Hughes Medical Institute Investigator when this work was initiated. We thank Eric Lander, Rudolf Jaenisch, Doeke Hekstra, Joseph Kirschvink for their helpful discussion and insights. We thank Leslie Gaffney for creating and editing figures.

---

**Extended Data Fig. 1.**
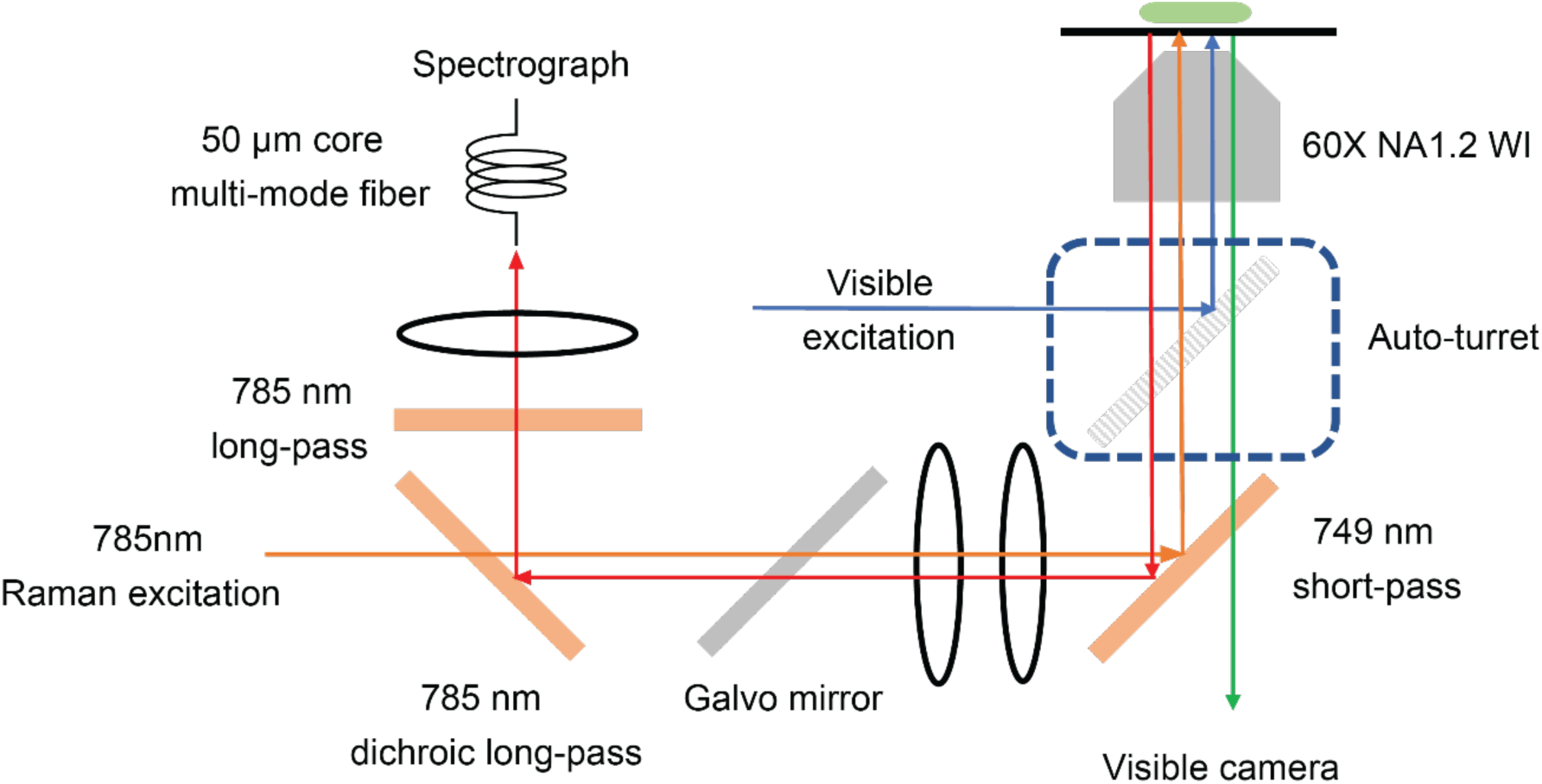
A multi-modal Raman microscope capable of fluorescence imaging and Raman microscopy. Schematic of a Raman microscope integrated with a wide-field fluorescence microscope for simultaneous detection of nuclei staining, bright field, fluorescence channels, and Raman images.

**Extended Data Fig. 2.**
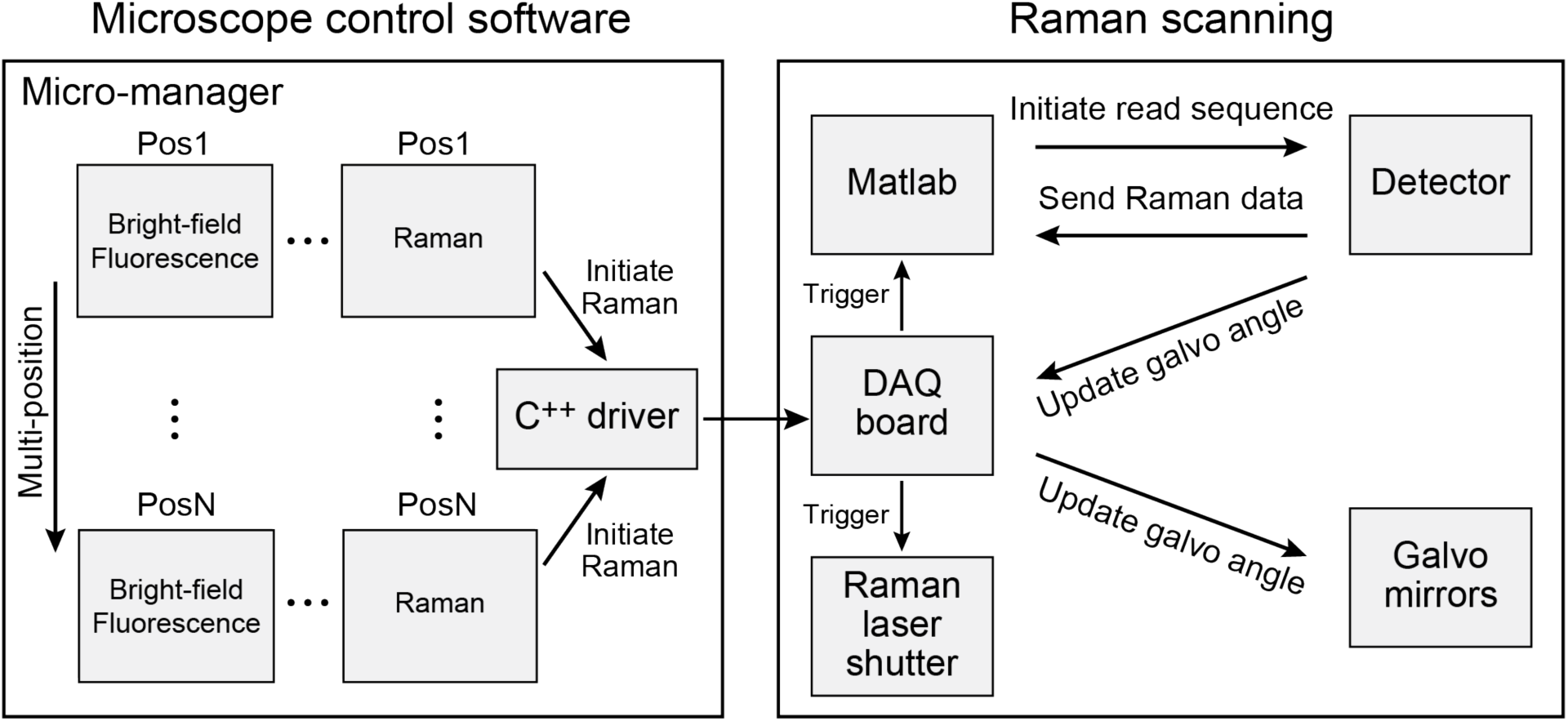
Overview of high-throughput Raman imaging software used in the study. A general-purpose microscope control software Micro-manager and a custom MATLAB script were combined to enable automated multi-modal measurements. Under Micro-manager, a Raman channel was registered as a ‘dummy’ channel along with brightfield and fluorescence channels. Micro-manager was responsible for changing the field of view (FOV) and imaging modality. During the Raman sequence, Micro-manager communicated with a digital acquisition (DAQ) board, through which a transistor-to-transistor logic (TTL) signal was generated to initiate the scanning sequence. Upon detection of the TTL signal, the MATLAB script controlled the Raman detector, laser shutter, and updated the galvo mirror angles through the DAQ board.

**Extended Data Fig. 3.**
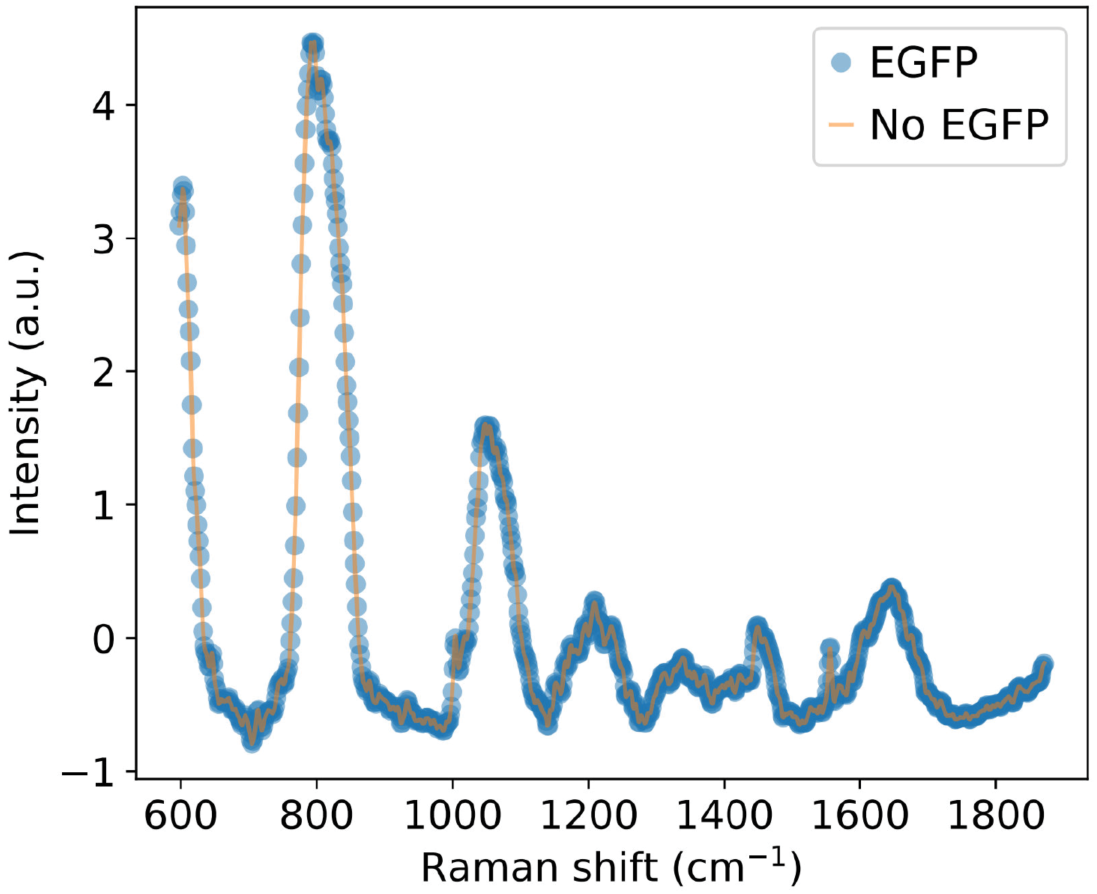
GFP does not interfere in Raman spectra measurement. Raman spectra of culture media with (blue) and without (orange) GFP at physiological concentration.

**Extended Data Fig. 4.**
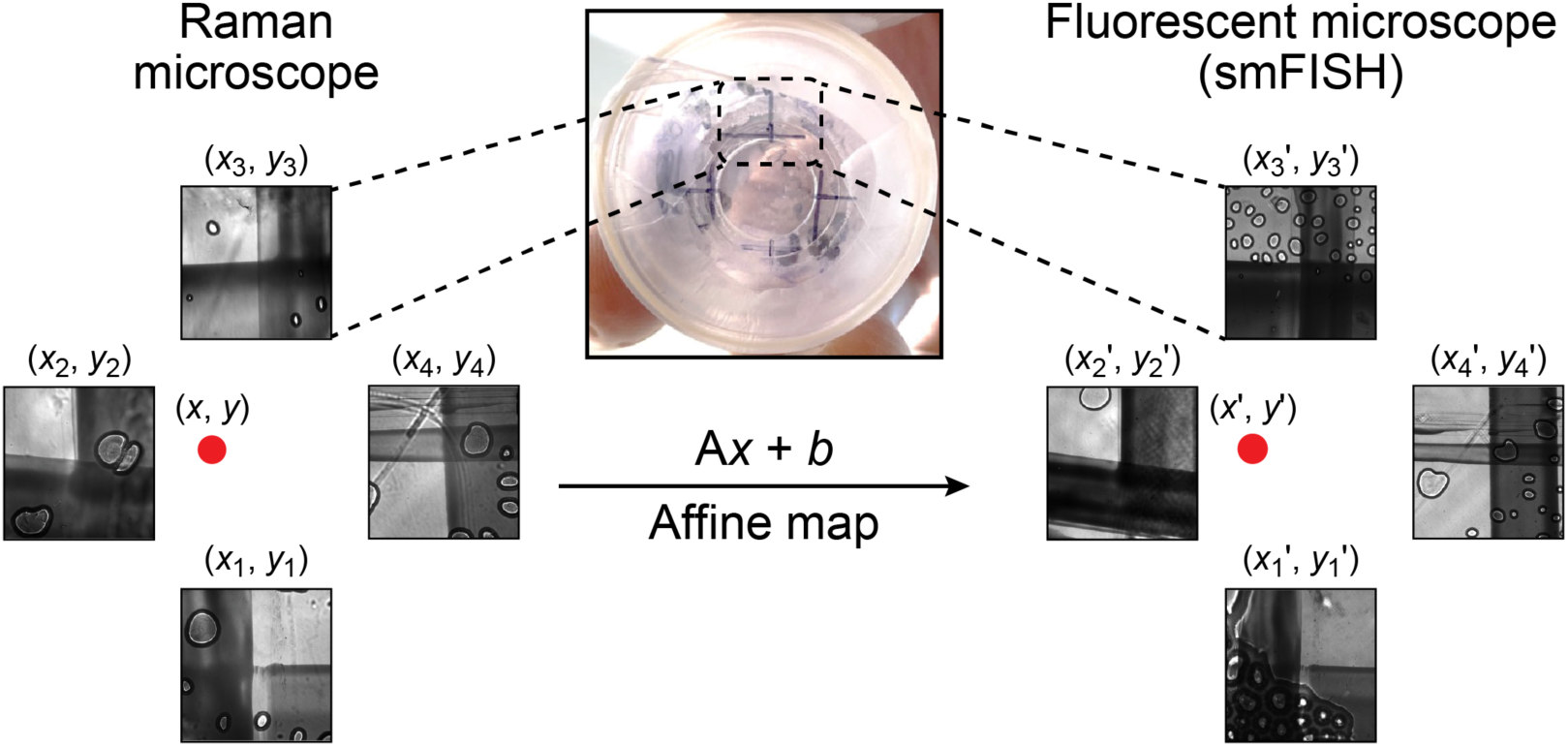
Image registration between the Raman and smFISH microscope using control points. Control points were inscribed under petri dishes with permanent markers and the coordinates were measured prior to any data acquisition. After Raman measurement and smFISH processing, samples were placed back to the microscope and control point coordinates were remeasured. Then, affine mapping was used to update the FOV coordinates to locate the exact same cells.

**Extended Data Fig. 5.**
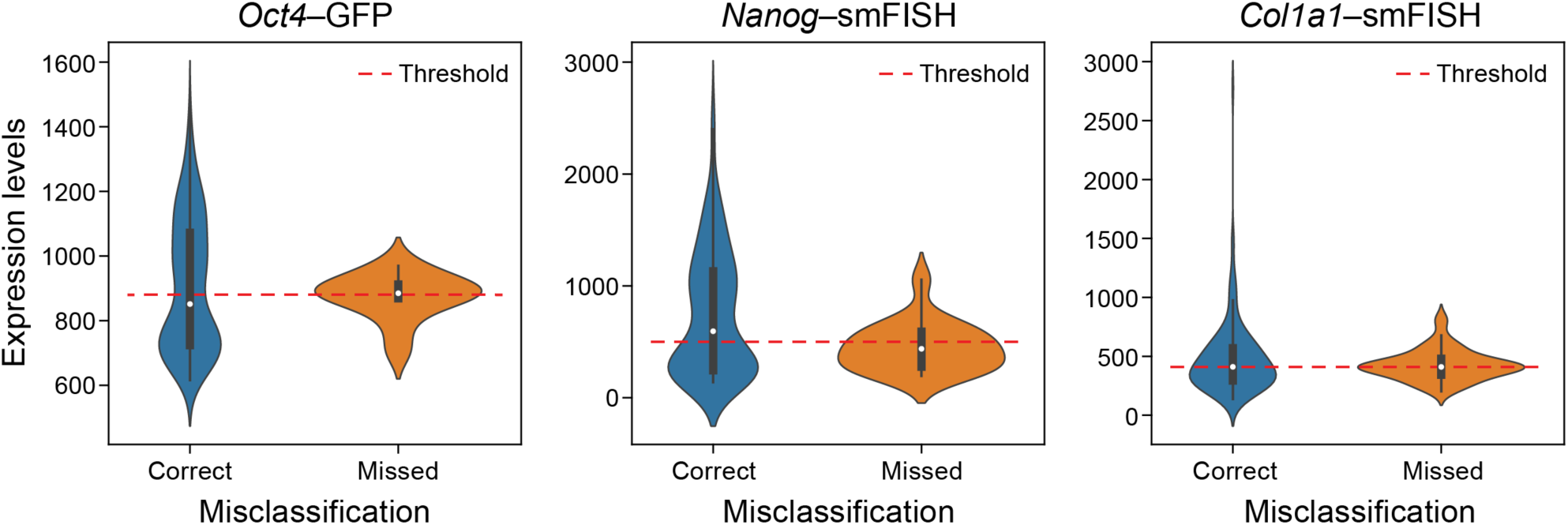
Misclassification of genes in the cell mixture classification experiment occurs when the ground truth smFISH is near the expression threshold. Distribution of measured smFISH expression level (y axis) for cells correctly (blue) or incorrectly (orange) classified by their Raman spectra for the expression of that gene. Horizontal line: an example threshold used for the logistic regression classifier.

**Extended Data Fig. 6.**
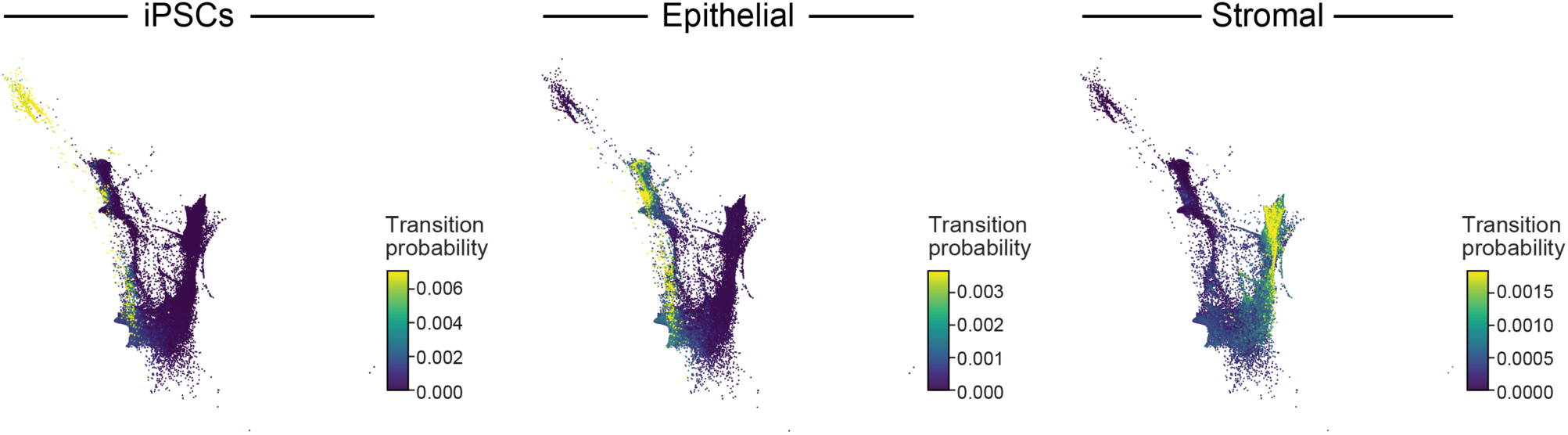
Cell transition probabilities inferred by Waddington-OT from scRNA-seq during reprogramming. Force-directed layout embedding (FLE) of scRNA-seq profiles (dots) from days 8 to 14.5 of reprogramming (dots) colored by the transition probability of each cell as inferred by Waddington-OT to be an ancestor of iPSCs (left), epithelial cells (middle) or stromal cells (right) at day 14.5.

**Extended Data Fig. 7.**
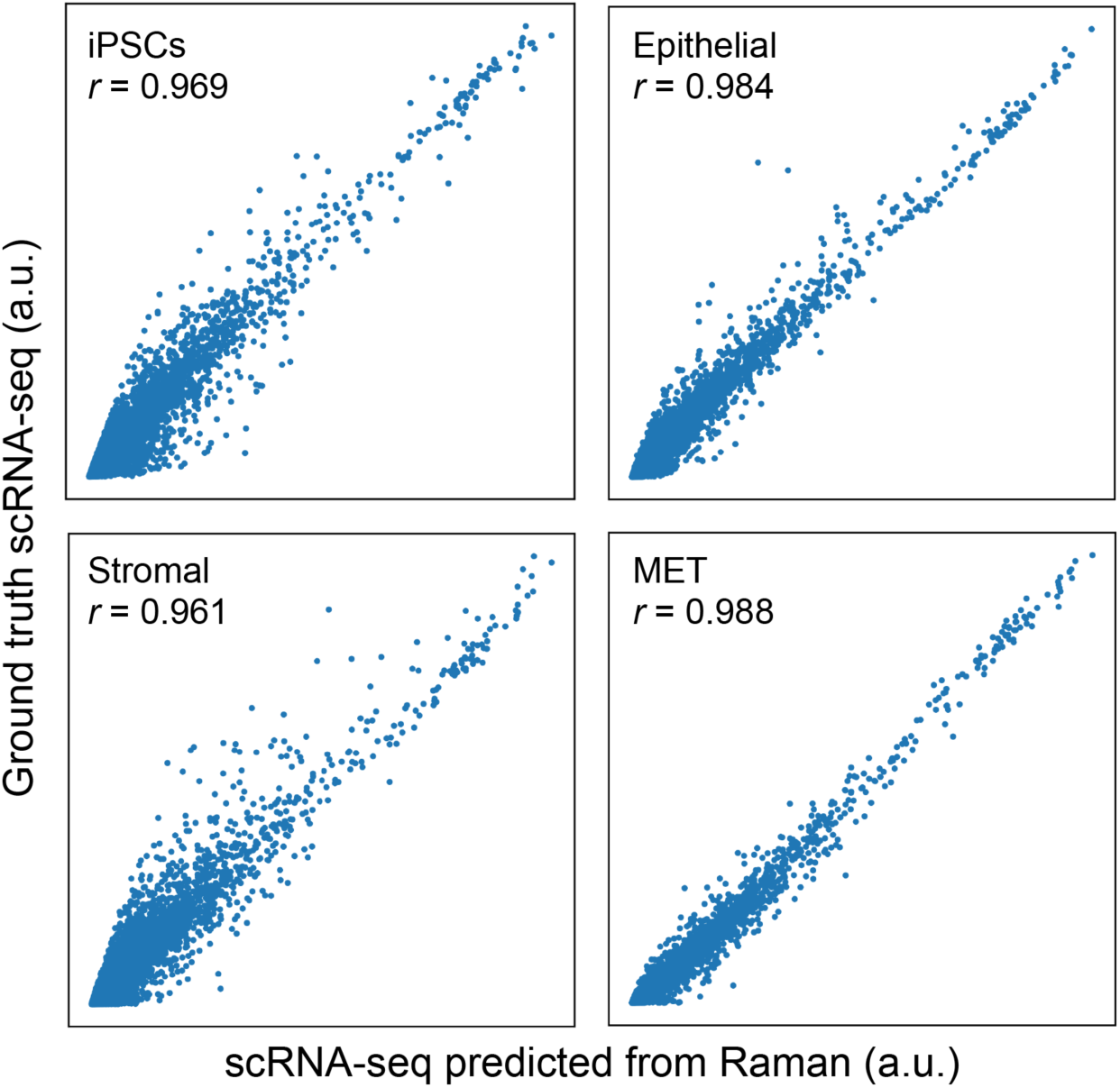
Raman-predicted and scRNA-seq measured pseudo-bulk profiles are well correlated across cell types. ScRNA-seq measured (y axis) and R2R-predicted (x axis) expression for each gene (dot) in pseudo-bulk RNA profiles averaged across cells labeled as iPSC (top left), epithelial (top right), stromal (bottom left) and MET (bottom right). Pearson’s r is denoted at the top left corner.

**Extended Data Fig. 8.**
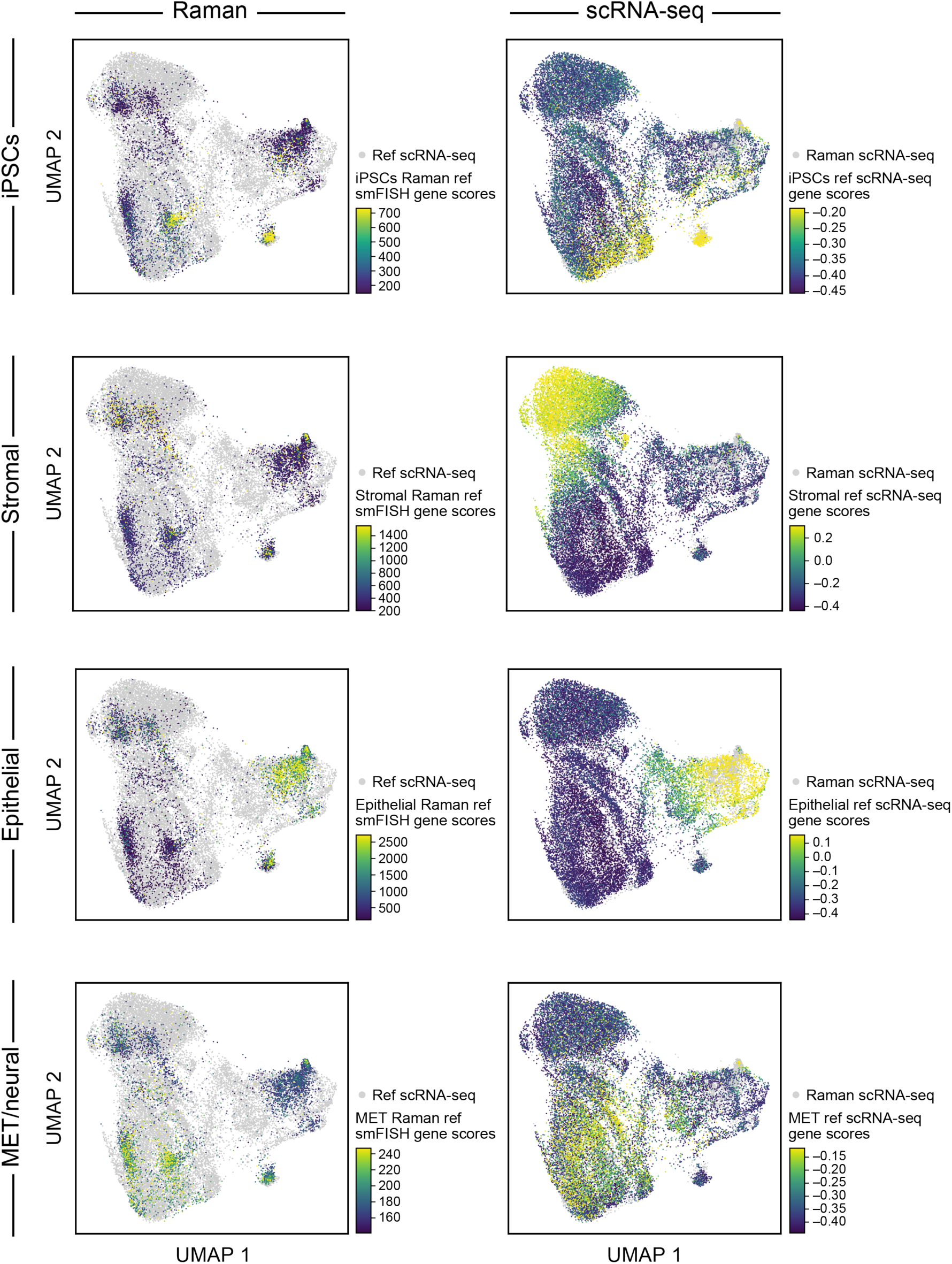
Measured and Raman-predicted single cell profiles co-embed well as reflected by gene scores for each cell type. UMAP co-embedding of Raman predicted RNA profiles and measured scRNA-seq profiles (dots) colored by scores of marker gene for different cell types (rows) determined by smFISH measurements (left, for cells with Raman-predicted profiles) or real scRNA-seq measurements (right, for cells with scRNA-seq profiles).

**Extended Data Fig. 9.**
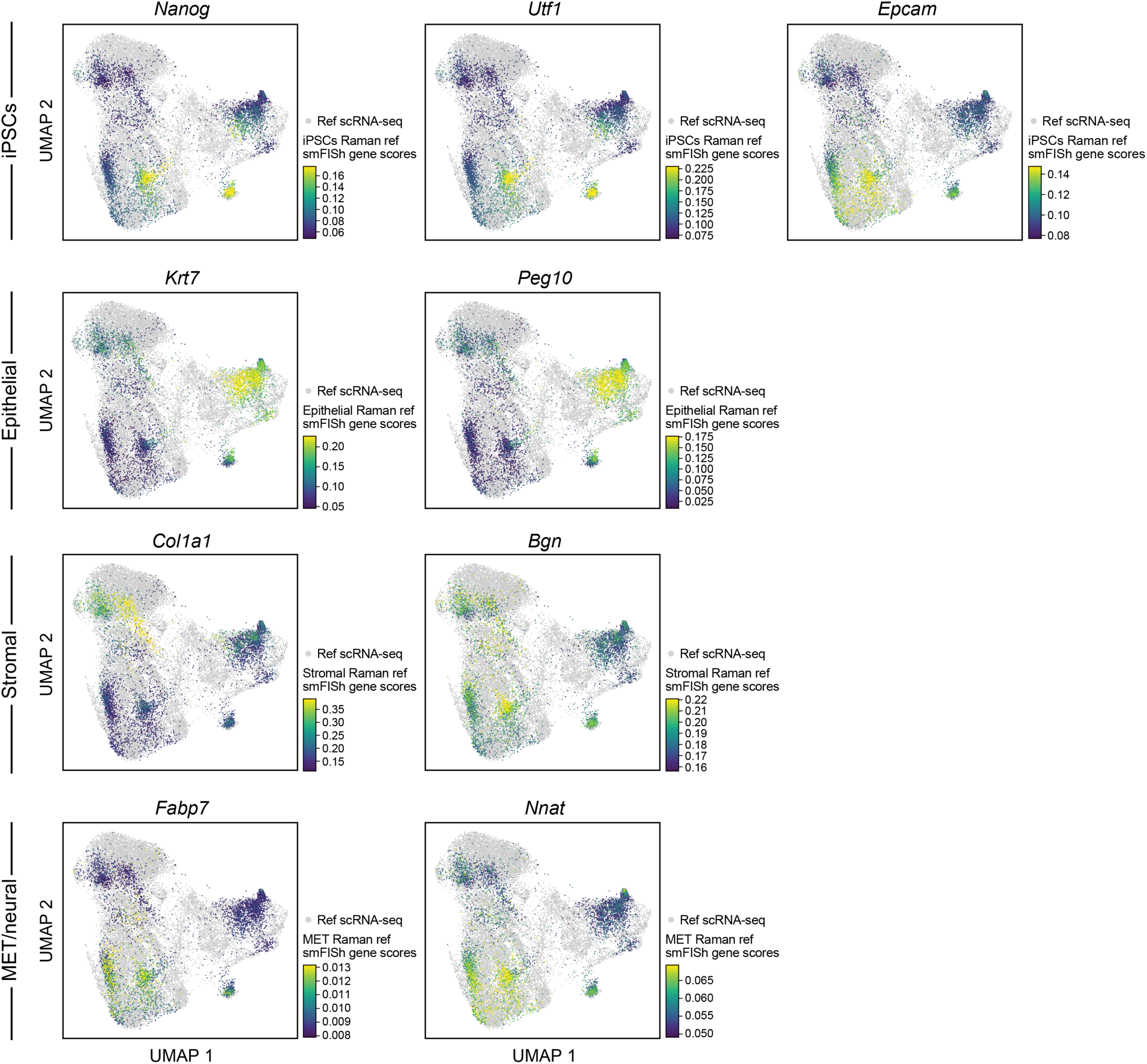
Measured and Raman-predicted single cell profiles co-embed well as reflected by smFISH measurement of Raman cells. UMAP co-embedding of Raman predicted RNA profiles and measured scRNA-seq profiles (dots) where the Raman cells are colored by smFISH measurement of each of nine anchor genes.

**Extended Data Fig. 10.**
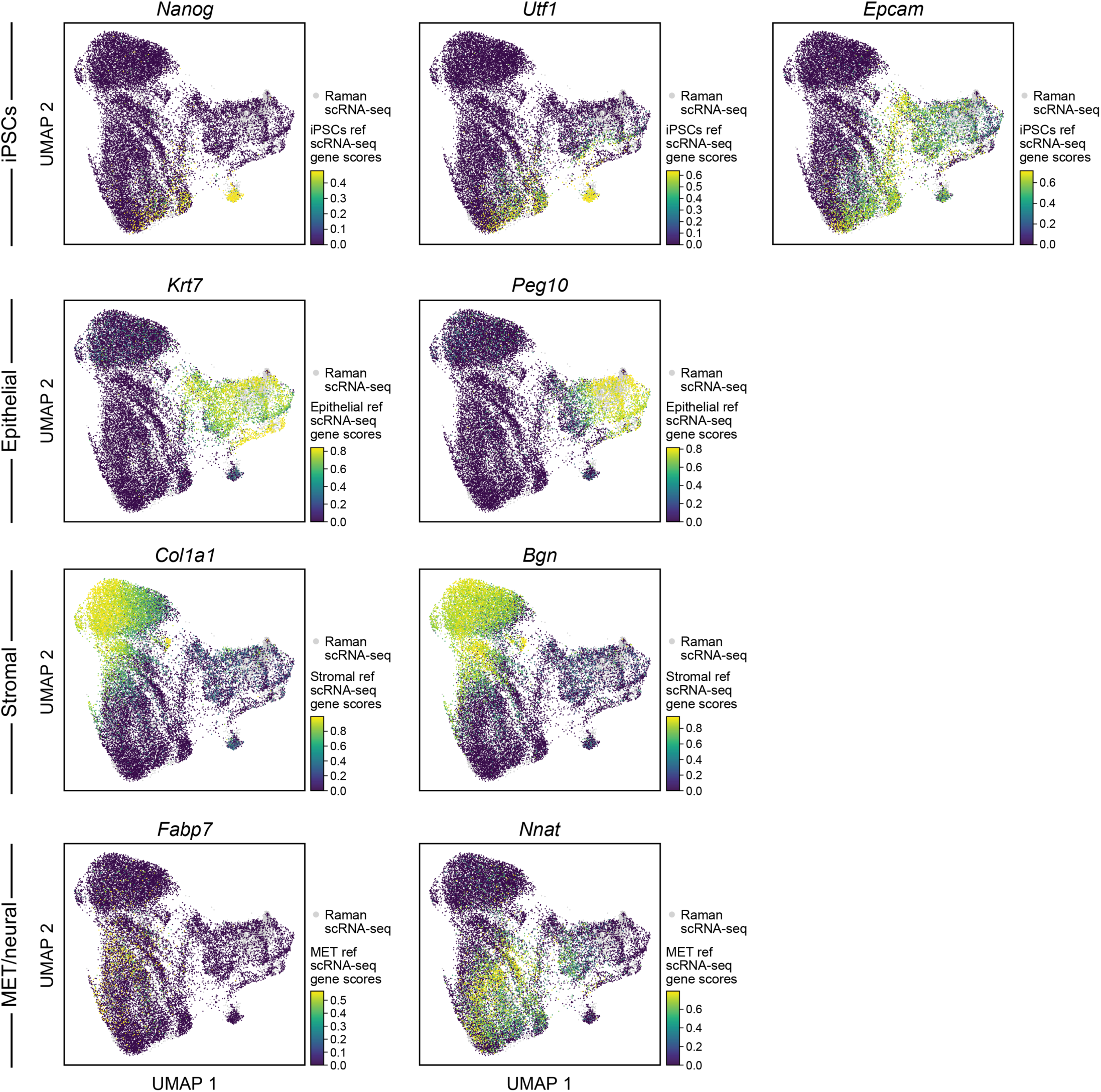
Measured and Raman-predicted single cell profiles co-embed well as reflected by scRNA-seq based expression of nine anchor genes. UMAP co-embedding of Raman predicted RNA profiles and measured scRNA-Seq profiles (dots) where the scRNA-seq profiled cells are colored by scRNA-seq measured expression of each of nine anchor genes.

**Extended Data Fig. 11.**
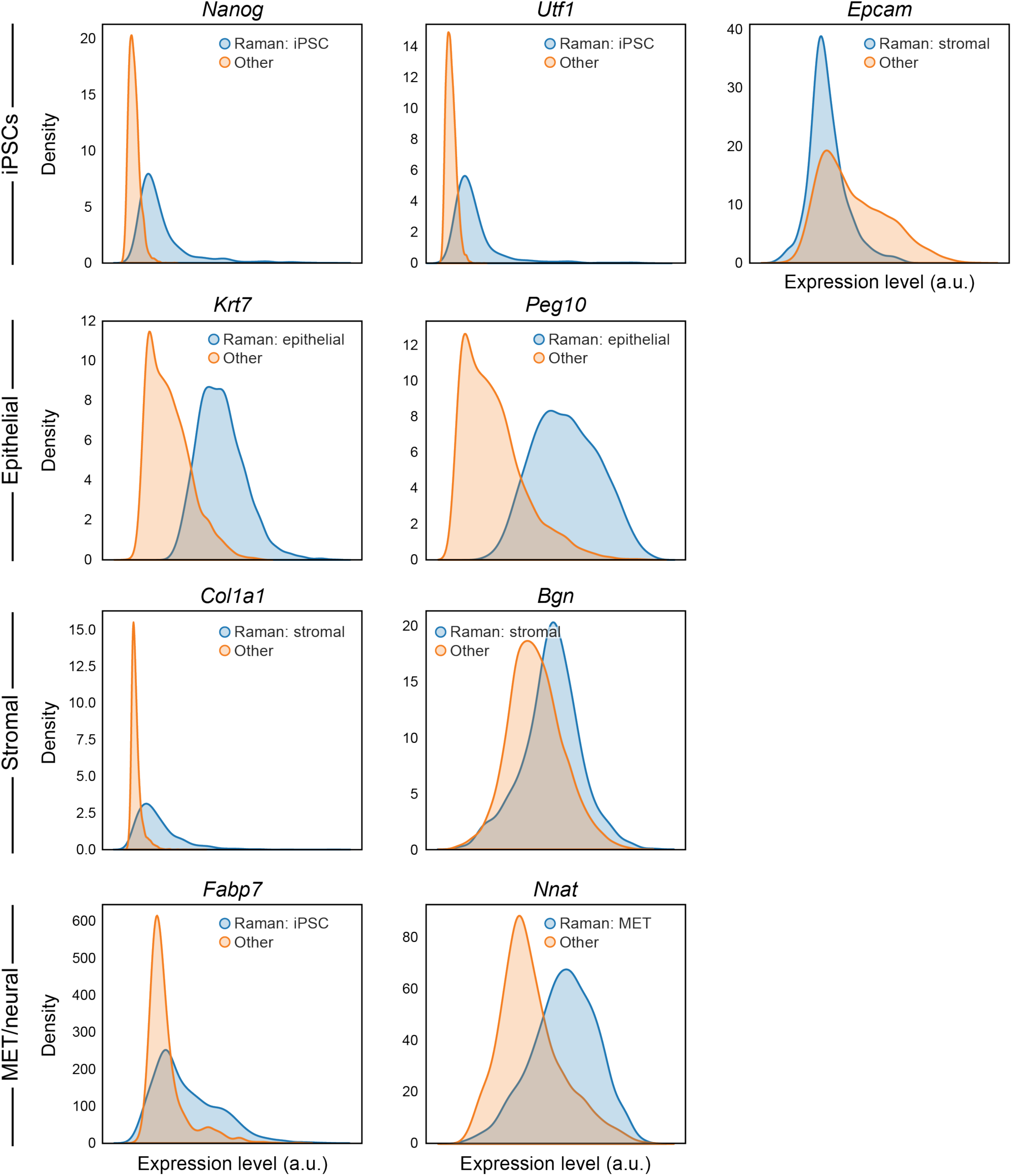
Distributions of expression of marker genes based on R2R-predicted profiles. Distributions (density plots) of the predicted expression in Raman2RNA inferred profiles for each marker gene (panel) in its expected corresponding cell type (blue, based on the predicted expression profiles) and all other cells (orange).

**Extended Data Fig. 12.**
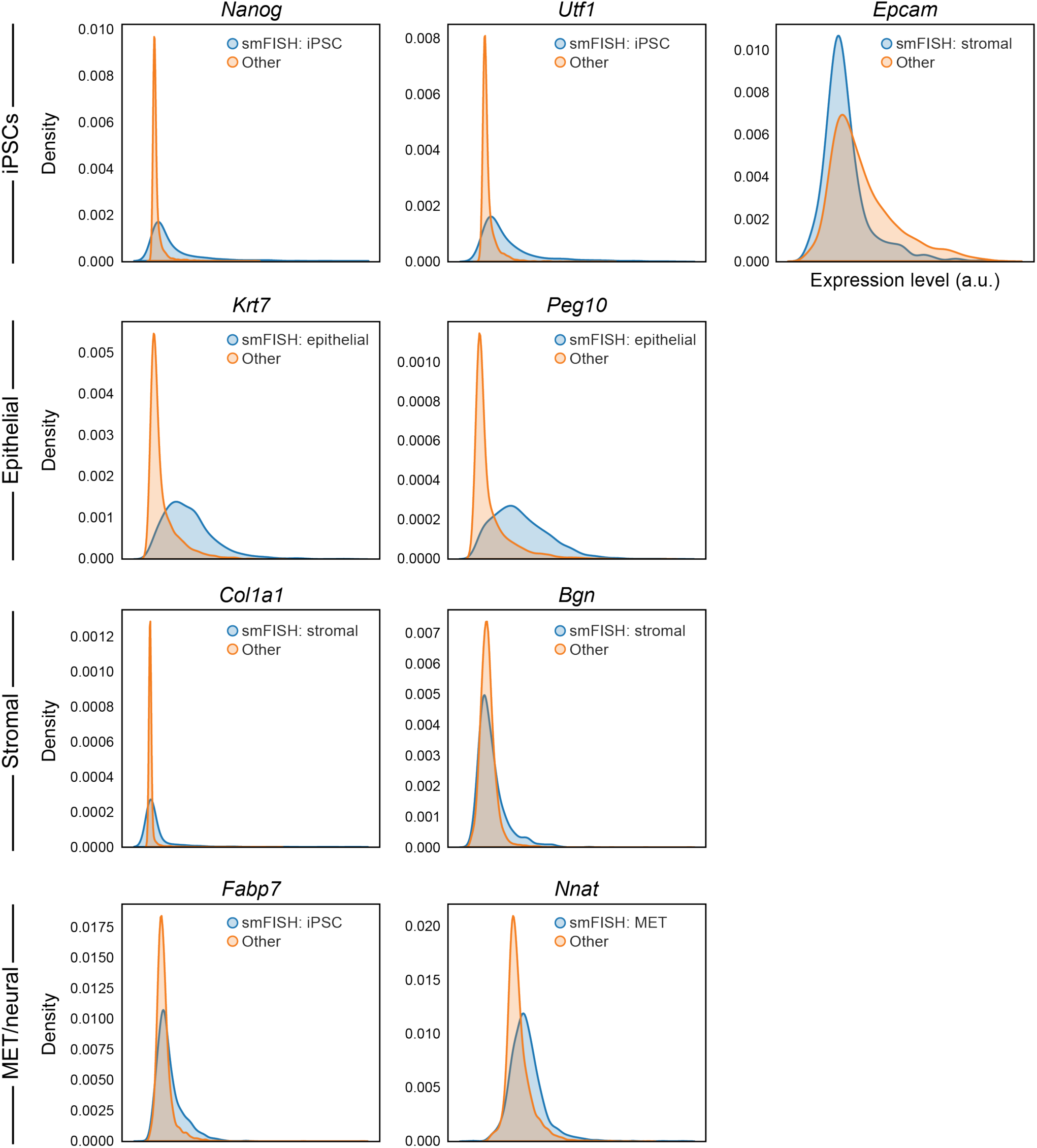
Distributions of expression of marker genes based on real smFISH profiles. Distributions (density plots) of the real smFISH profiles for each marker gene (panel) in its expected corresponding cell type (blue, based on the R2R *predicted* expression profiles) and all other cells (orange).

**Extended Data Fig. 13.**
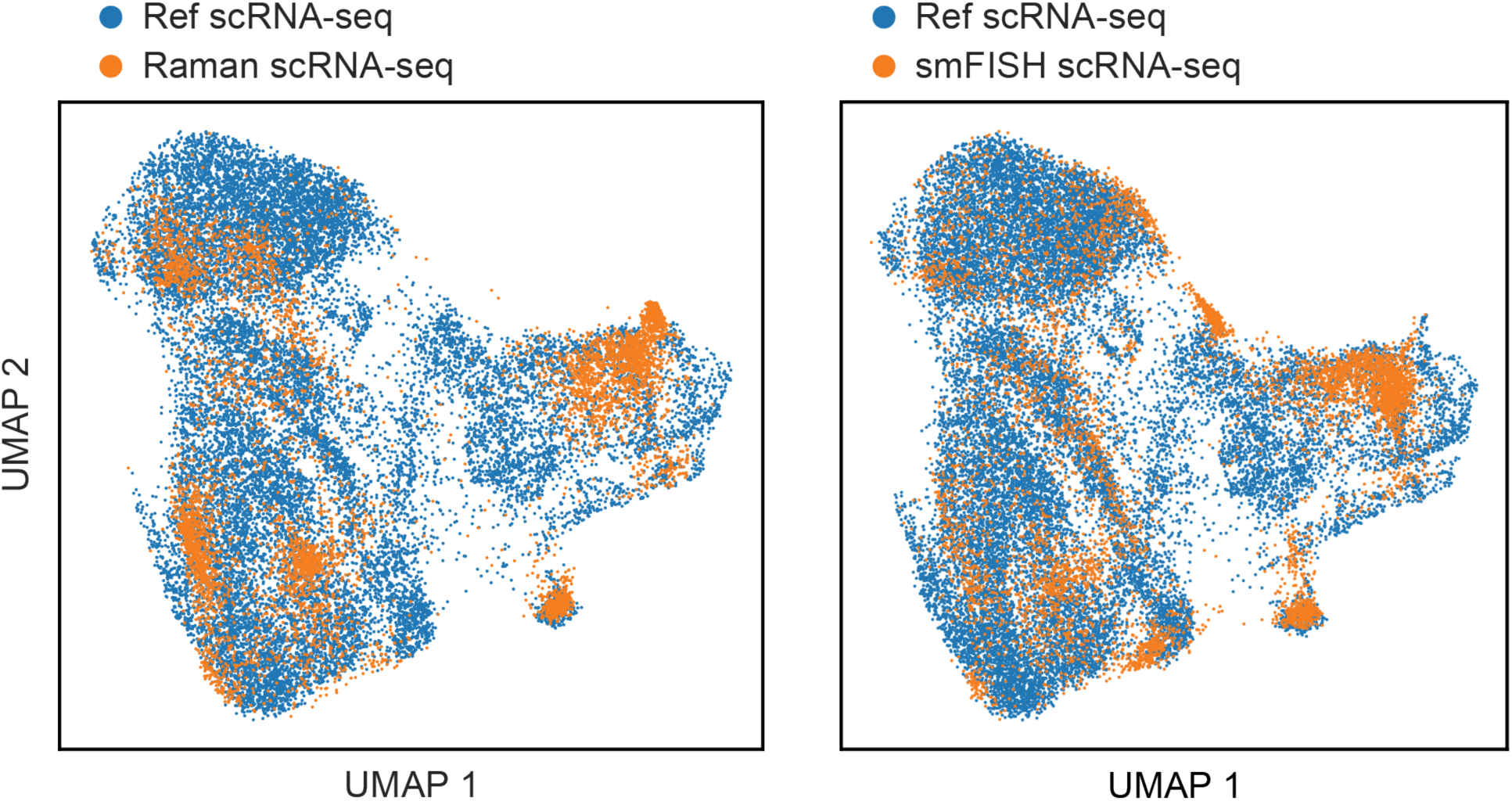
RNA profiles predicted directly from 9 anchor smFISH measurements lead to reduced variance compared to scRNA-seq. UMAP co-embedding of cells from scRNA-seq (blue) and Raman (orange) experiments, with the latter based on either the Raman-predicted RNA profiles (left) or only smFISH-predicted RNA profiles (right).

**Extended Data Fig. 14.**
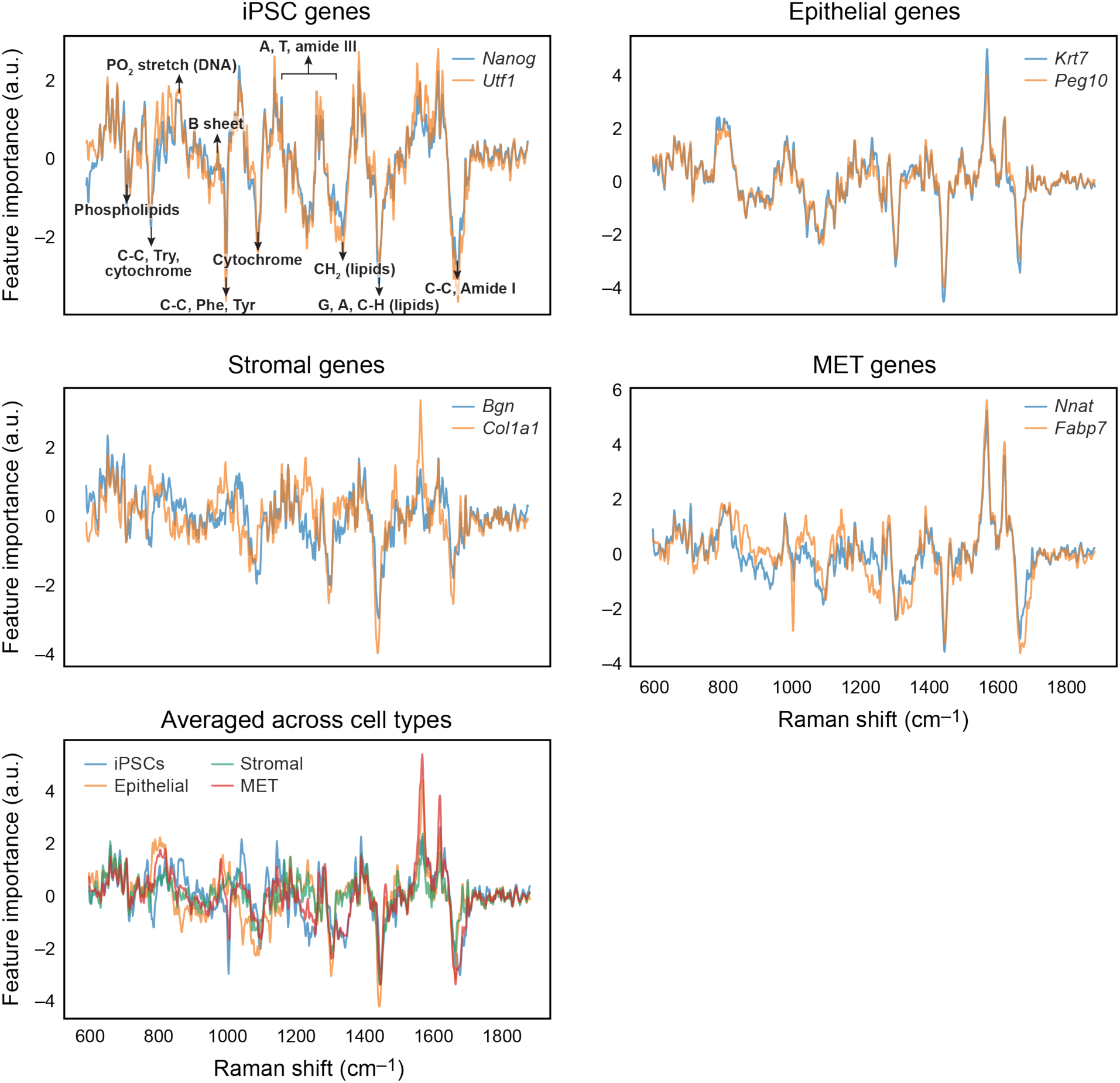
Raman spectral feature importance scores for each smFISH anchor gene and its average across all genes for a cell type. Feature importance scores (y axis) for marker genes of each cell type (top two rows), and for all cell types (bottom row), along the Raman spectrum (x axis). Known signals^18^ are annotated in the top left panel (identical to Fig. 3k).

**Extended Data Fig. 15.**
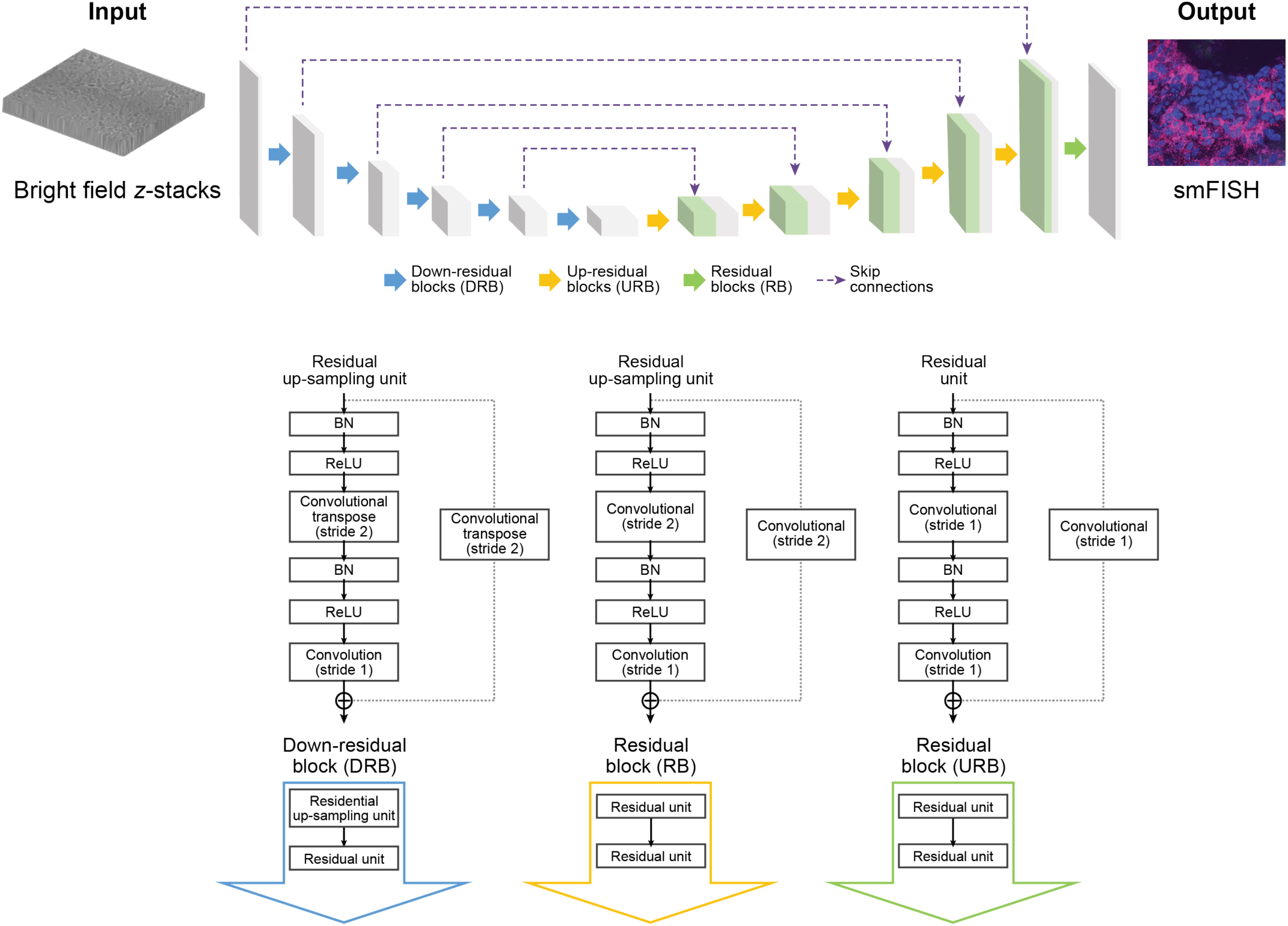
Neural network-based prediction of smFISH using brightfield z-stacks.

